# Multivoxel codes for representing and integrating acoustic features in human cortex

**DOI:** 10.1101/730234

**Authors:** Ediz Sohoglu, Sukhbinder Kumar, Maria Chait, Timothy D. Griffiths

**Author notes:** MRC Cognition and Brain Sciences Unit, University of Cambridge, 15 Chaucer Road, Cambridge, CB2 7EF, United Kingdom. **Address correspondence to:** Ediz Sohoglu.

## Abstract

Using fMRI and multivariate pattern analysis, we determined whether acoustic features are represented by independent or integrated neural codes in human cortex. Male and female listeners heard band-pass noise varying simultaneously in spectral (frequency) and temporal (amplitude-modulation [AM] rate) features. In the superior temporal plane, changes in multivoxel activity due to frequency were largely invariant with respect to AM rate (and vice versa), consistent with an independent representation. In contrast, in posterior parietal cortex, neural representation was exclusively integrated and tuned to specific conjunctions of frequency and AM features. Direct between-region comparisons show that whereas independent coding of frequency and AM weakened with increasing levels of the hierarchy, integrated coding strengthened at the transition between non-core and parietal cortex. Our findings support the notion that primary auditory cortex can represent component acoustic features in an independent fashion and suggest a role for parietal cortex in feature integration and the structuring of acoustic input.

**Significance statement:** A major goal for neuroscience is discovering the sensory features to which the brain is tuned and how those features are integrated into cohesive perception. We used whole-brain human fMRI and a statistical modeling approach to quantify the extent to which sound features are represented separately or in an integrated fashion in cortical activity patterns. We show that frequency and AM rate, two acoustic features that are fundamental to characterizing biological important sounds such as speech, are represented separately in primary auditory cortex but in an integrated fashion in parietal cortex. These findings suggest that representations in primary auditory cortex can be simpler than previously thought and also implicate a role for parietal cortex in integrating features for coherent perception.

## Introduction

In structuring the auditory scene, the brain must carry out two fundamental computations. First, it must derive *independent* representations of component acoustic features so that task-relevant features can be prioritized and task-irrelevant ones ignored. Second, to solve the well-known “binding problem”, the brain must subsequently *integrate* these separated representations into a coherent whole so that the features of a relevant sound source can be tracked successfully in cluttered scenes. Whether representations of stimulus features are independent or integrated is a longstanding issue in psychology (Treisman and Gelade, 1980; Ashby and Townsend, 1986) and neuroscience (Di Lollo, 2012; Soto et al., 2018). Even when not explicitly framed using these terms, many questions concerning sensory systems can be formalized in terms of representational independence versus integration (Soto et al., 2018).

It is widely believed that auditory processing is hierarchically organized and that neural representations are progressively transformed from independent to integrated codes as sensory information ascends the auditory pathway (Rauschecker and Tian, 2000; Bizley and Cohen, 2013). Thus, while neurons in low-level regions might respond to single stimulus features, higher-level neurons should show more complex tuning properties and respond to conjunctions of features. Precisely where along this continuum human primary auditory cortex (and regions beyond) fit within this conception of the auditory system has been the subject of debate.

Based on presumed similarities with the visual system, early models proposed that representations in primary auditory cortex were primarily independent, instantiated as topographically organized “feature maps” (see Nelken et al., 2003). According to such accounts, the integration of features is a computation that should most reliably be observed in non-primary regions. However, animal physiology studies demonstrate highly non-linear neural responses already at the level of primary auditory cortex, suggestive of an integrated coding scheme (deCharms et al., 1998; Nelken et al., 2003; Chi et al., 2005; Wang et al., 2005; Christianson et al., 2008; Atencio et al., 2009; Bizley et al., 2009; Sadagopan and Wang, 2009; Sloas et al., 2016). The extent to which this also applies in humans remains unclear. While there are many sources of human imaging evidence that are potentially relevant to this issue, particularly investigations of how low-level acoustic features and higher-level categories are represented in cortical activity (Davis and Johnsrude, 2003; Zatorre et al., 2004; Cusack, 2005; Kumar et al., 2007; Staeren et al., 2009; Leaver and Rauschecker, 2010; Teki et al., 2011; Giordano et al., 2013; Norman-Haignere et al., 2015; Overath et al., 2015; Allen et al., 2017), fewer studies have directly tested and quantified the extent of representational independence versus integration in human cortex.

In the current study, we used fMRI and multivariate pattern analysis to determine the extent to which acoustic features are represented by independent or integrated multivoxel codes and how those codes are expressed over the human cortical hierarchy. In general, multivariate approaches allow sensory features to be more directly linked to their representation in neural response patterns (Tong and Pratte, 2012; Kriegeskorte and Kievit, 2013; Haynes, 2015), in contrast to traditional univariate analysis of overall regional differences in signal amplitude. In this study, an approach based on MANOVA (Allefeld and Haynes, 2014) allowed us to estimate the contribution of single acoustic features to the observed multivoxel patterns (reflecting independent coding), as opposed to non-linear interactions between the features that may arise at the level of object perception (integrated coding). Moreover, by acquiring whole-brain fMRI, we were able to characterize neural representations simultaneously across the entire human cortex, in contrast to more localized physiological recordings in animals.

Participants listened to band-pass noise varying simultaneously in frequency (a spectrally-based feature) and amplitude modulation (AM) rate (temporally-based; see Figure 1A). We chose to investigate these two acoustic features as they are sufficient alone to characterize much of the information present in biologically important sounds such as speech (Shannon et al., 1995; Roberts et al., 2011).

**Figure 1.**
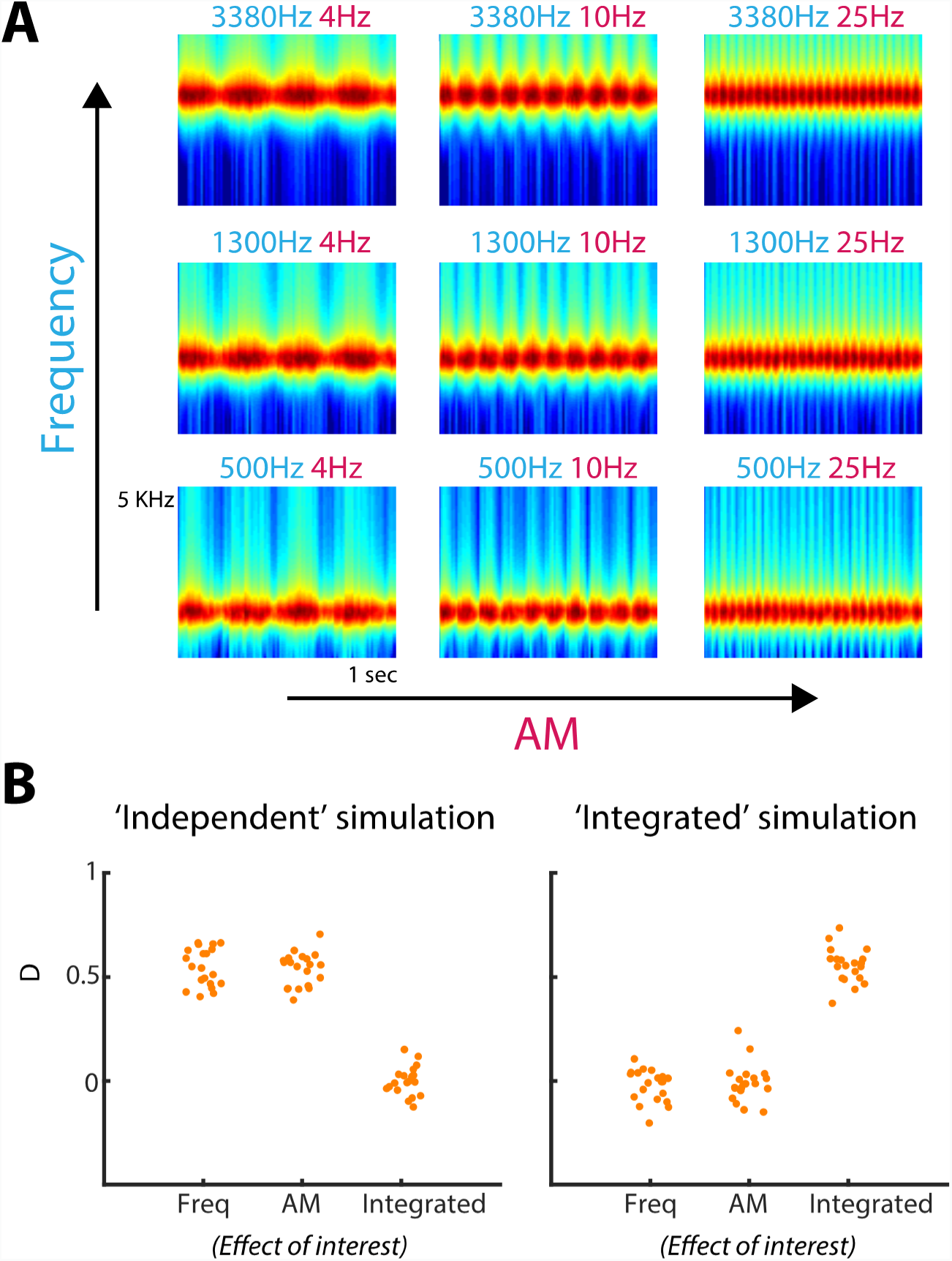
A) Spectrograms of the nine stimuli, equally spaced on a scale of ERB-rate (Moore and Glasberg, 1983) and smoothed to obtain a temporal resolution similar to the Equivalent Rectangular Duration (Plack and Moore, 1990). The cyan- and magenta-colored text above each spectrogram indicate the center carrier frequency and AM rate of the bandpass noise, respectively. B) Multivariate pattern distinctness estimates for each effect of interest, when activity patterns were simulated using an independent representation (left-side graph) or an integrated representation (right-side graph). Each data point represents the pattern distinctness for a single iteration (“participant”) of the simulation. Freq, Frequency. D, Pattern distinctness.

## Methods

### Participants

Twenty participants (eleven female), aged between 18 and 27 years (mean = 23, SD = 2.4), were tested after being informed of the study’s procedure, which was approved by the research ethics committee of University College London. All reported normal hearing, normal or corrected-to-normal vision, and had no history of neurological disorders.

### Stimuli

The stimulus consisted of narrow (third of an octave) bandpass noise, amplitude modulated sinusoidally with 80% depth (see Figure 1A). Each sound was presented for one second and varied across trials in center carrier frequency (from hereon, “frequency”) and amplitude modulation rate (“AM”). Frequency (500, 1300 and 3380 Hz) and AM (4, 10 and 25 Hz) were equally spaced on a logarithmic scale. Importantly for the purpose of assessing independent and integrated feature coding (see First-level statistics section below), frequency and AM varied simultaneously and in an orthogonal fashion, such that every frequency was paired with every AM (i.e. nine stimuli in total, arranged as a 3 x 3 factorial design). The relatively slow AM rates precluded the perception of pitch associated with the temporal modulation. In addition, the carrier center frequencies and bandwidths were chosen to avoid detectable spectral cues from resolved sidebands in the stimulus (Moore, 2003).

Stimuli were matched in terms of their RMS amplitude and shaped with 20 ms raised-cosine onset and offset ramps. Bandpass noise was synthesized independently on each presentation (with a sampling rate of 44100 Hz) and delivered diotically through MRI-compatible insert earphones (S14, Sensimetrics Corporation). To compensate for resonances in the frequency response of the earphones, the stimuli were digitally preprocessed using the filters and software provided with the earphones.

### Procedure

Stimulus delivery was controlled with Cogent toolbox (http://www.vislab.ucl.ac.uk/cogent) in Matlab (MathWorks). Participants were scanned for five runs, each lasting around ten minutes consisting of sixteen repetitions of the nine stimuli. For one participant, there was insufficient time to scan for the fifth run because of technical difficulties. Stimuli were grouped into blocks of eighteen sounds within which all nine stimuli appeared twice and in random order. The inter-stimulus interval ranged uniformly between 2000 and 4000 ms.

Participants were instructed to listen carefully to the sounds while looking at a central fixation cross and press a button (with their right hand) each time a brief (150 ms duration) white-noise interruption occurred during sound presentation. These white-noise interruptions were unmodulated in their amplitude profile and occurred on a small percentage (~6%) of stimuli (once every block of eighteen sounds). Group performance was near ceiling, confirming engagement with the task. The average hit rate was. 98 (ranging from. 8 to 1 across participants; SEM = .014) with no false alarms.

To estimate the perceived saliency of the sounds, two participants from the main fMRI experiment and four new participants (two female; mean age = 29 years, SD = 4) completed a short behavioral session similar in procedure to Petsas et al. (2016). These participants listened to all pairwise combinations of the nine sounds (eight pairs for each of the nine sounds; separated by 200 ms of silence) and were asked to judge on each trial which of the two sounds was more salient. Pairs were presented three times in random order, with the order of the sounds within a pair counterbalanced across trials.

To estimate perceived loudness, we used the loudness model of Moore et al. (2016), as implemented in Matlab (http://hearing.psychol.cam.ac.uk/TVLBIN/tv2016Matlab.zip). As the model output differs slightly for different noise samples of the same condition, we generated an entire (single-participant) stimulus set in the same way as was done for the main experiment and submitted each stimulus to the model. We computed the time-varying long-term loudness, averaged over the duration of the stimulus and across noise samples within each of the nine stimuli.

### Image acquisition

Imaging data were collected on a Siemens 3 Tesla Quattro MRI scanner (http://www.siemens.com) at the Wellcome Trust Centre for Human NeuroImaging, University College London. A total of 175 echo planar imaging (EPI) volumes were acquired per run, using a 32-channel head coil and continuous sequence (TR = 3.36 sec; TE = 30 ms; 48 slices covering the whole brain; 3 mm isotropic resolution; matrix size = 64 x 74; echo spacing = 0.5 ms; orientation = transverse). After the third run, field maps were acquired (short TE = 10 ms; long TE = 12.46 ms). During the functional scans, we also obtained physiological measures of each participant’s breathing and cardiac pulse. Because of technical issues, physiological measures were not available for two participants. The experimental session concluded with the acquisition of a high-resolution (1 x 1 x 1 mm) T1-weighted structural MRI scan.

### Image processing

fMRI analysis was performed in SPM12 (http://www.fil.ion.ucl.ac.uk/spm). After discarding the first three volumes to allow for magnetic saturation effects, the remaining images were realigned and unwarped to the first volume to correct for movement of participants during scanning. Also at the unwarping stage, the acquired field maps were used to correct for geometric distortions in the EPI due to magnetic field variations. Realigned images were co-registered to the mean functional image and then subjected to multivariate statistical analysis, generating searchlight maps from unsmoothed data in each participant’s native space (see First-level statistics section below). Searchlight maps were subsequently normalized to the Montreal Neurological Institute (MNI) template image using the parameters from the segmentation of the structural image (resampled resolution: 2 x 2 x 2 mm) and smoothed with a Gaussian kernel of 6 mm full-width at half-maximum. Where additional univariate analyses are reported, realigned images were spatially normalized and smoothed first before statistical analysis.

### First-level statistics

Statistical analysis was based on the general linear model (GLM) of each participant’s fMRI time series, using a 1/128 Hz highpass filter and AR1 correction for auto-correlation. The design matrix comprised the auditory stimulus events, each modeled as a stick (delta) function and convolved with the canonical haemodynamic response function. Separate columns were specified for each of the nine stimuli, in addition to a column for target sounds (to remove variance associated with the white noise interruptions and the button presses). Additional columns were specified for the six movement parameters and the mean of each run. Cardiac and respiratory phase (including their aliased harmonics) as well as heart rate and respiratory volume were modeled using an in-house Matlab toolbox (Hutton et al., 2011). This resulted in fourteen physiological regressors in total: six each for cardiac and respiratory phase and one each for heart rate and respiratory volume.

For statistical inference, we used cross-validated multivariate analysis of variance (Allefeld and Haynes, 2014), as implemented in the cvMANOVA toolbox in Matlab (version 3; https://github.com/allefeld/cvmanova). For each participant this method measures the pattern distinctness D, a cross-validated version of one of the standard multivariate statistics: Lawley-Hotelling’s trace. D quantifies the multivoxel variation in activity attributable to an experimental contrast, relative to unexplained variation or noise (for examples of previous applications, see Guggenmos et al., 2016; Christophel et al., 2017, 2018; Dijkstra et al., 2017). Thus, D is the multivariate extension of the univariate F-statistic in ANOVA and is a clearly interpretable measure of effect size. This is in contrast to classification accuracy from pattern decoders, which is dependent on the particular algorithm used as well as the amount of data and partitioning into training and test sets (see Allefeld and Haynes, 2014). Cross-validation ensures that the expected value of D is zero if two voxel patterns are not statistically different from each other, making D a suitable summary statistic for group-level inference (e.g. with the one-sample t-test). Note that because of this cross-validation, D can sometimes be negative if its true value is close to zero in the presence of noise.

When applied to the simple case of only two stimuli, the pattern distinctness D is a measure of between-stimulus pattern dissimilarity and is closely related to the (cross-validated) Mahalanobis distance, which is argued to be a more reliable and accurate metric for characterizing multivoxel patterns than the correlation or Euclidean distance (Kriegeskorte et al., 2006; Ejaz et al., 2015; Walther et al., 2016). Like the Mahalanobis distance, D takes into account the spatial structure of the noise (GLM residuals) by normalizing the multivoxel variation for an experimental effect by the noise covariance between voxels. As D is obtained from the GLM, cvMANOVA can also be used to test more complex contrasts such as main effects and interactions with a factorial design. As explained below, we can use such factorial contrasts to distinguish between independent and integrated neural coding.

cvMANOVA was performed as a searchlight analysis (Kriegeskorte et al., 2006) using spheres with a radius of three voxels (~9 mm; ~123 voxels of 3 x 3 x 3 mm) and constrained to voxels within the whole-brain mask generated by SPM during model estimation. Thus, for each participant and effect of interest, a whole-brain searchlight image was generated in which each voxel expressed the pattern distinctness D over that voxel and the surrounding neighborhood. As recommended by Allefeld and Haynes (Allefeld and Haynes, 2014), to correct for searchlight spheres near the brain mask boundaries containing fewer voxels, the estimate of D at each voxel was standardized by dividing by the square root of the number of voxels within the searchlight.

We tested the extent to which frequency and AM features are represented by independent or integrated neural codes by examining three effects of interest. If frequency and AM features are represented in an integrated fashion, then changes in these two features should combine non-linearly (non-additively) to influence multivoxel activity patterns (see Kornysheva and Diedrichsen, 2014; Erez et al., 2015). In other words, the effect of frequency should differ depending on AM (and vice versa). Thus, the first effect of interest was the interaction between frequency and AM and quantified the extent of integrated coding. If on the other hand, frequency and AM features are coded independently, then changes in these two features should result in a linear (additive) effect on activity patterns. An independent effect implies that changes in voxel patterns attributable to the frequency feature remain invariant with respect to AM (and vice versa): there is no interaction. Within the cvMANOVA framework, the extent of independence can be quantified by subtracting the interaction from the main effects (following equation 19 in Allefeld and Haynes 2014), resulting in the two other effects of interest: Independent coding of frequency and Independent coding of AM. These measures of independent coding are equivalent to those obtained from “cross-decoding” in classifier-based multivoxel pattern analysis (Formisano et al., 2008; Allefeld and Haynes, 2014; Kornysheva and Diedrichsen, 2014; Simanova et al., 2014).

Computational simulations confirm that the above effects of interest can successfully detect the presence of independent and integrated representations. For each of twenty “participants”, five “runs” and nine stimuli, we generated synthetic activity patterns over 123 voxels consisting of the true underlying pattern (normal random vector) added to some noise (signal-to-noise ratio set to 0.1). Within one run, there were sixteen repetitions of the nine stimuli. These synthetic data were then submitted to cvMANOVA resulting in a pattern distinctness estimate for each participant and effect of interest.

Two versions of the simulation were run, following Kornysheva and Diedrichsen (2014). In the first version, frequency and AM features were represented independently. That is, voxel patterns were generated separately for the two features and summed together to obtain voxel patterns (Y) for each of the nine stimuli with carrier center frequency f and AM rate m:

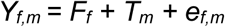

where F and T denote, respectively, the voxel pattern representations for the frequency and AM features and e the noise.

In the second version, frequency and AM were represented in an integrated fashion by generating a unique pattern for each of the nine stimuli. Thus, in this version of the simulation, the representation of frequency is inseparable from that of AM:

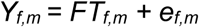

Here FT denotes the true pattern that was generated uniquely for each condition. In both versions, the resulting patterns were scaled to have the same mean and variance.

As Figure 1B shows, when frequency and AM were simulated as independent representations, the pattern distinctness D was significantly greater than zero when testing the independent (but not integrated) coding effects of interest (frequency: t(19) = 29.2, p < .001; AM: t(19) = 35.1, p < .001; Integrated: t(19) = −.104, p = .541). In contrast, when frequency and AM were represented in an integrated fashion, the reverse was true with a significant effect of integrated (but not independent) coding (frequency: t(19) = −1.39, p = .910; AM: t(19) = −.429, p = .664; Integrated: t(19) = 33.0, p < .001).

### Group-level statistics

Searchlight images were submitted to a group-level one-sample t-test under minimal assumptions using a nonparametric permutation procedure, as implemented in SnPM (http://warwick.ac.uk/snpm). We used 5000 iterations with 6 mm of variance smoothing (Nichols and Holmes, 2002) and constrained the analysis to voxels within the cortex (as defined by the probabilistic Harvard-Oxford cortical mask thresholded at 25%, distributed with FslView https://fsl.fmrib.ox.ac.uk). Statistical maps were thresholded voxelwise at p < .005 and clusterwise at p < .05 (familywise error [FWE] corrected for multiple comparisons).

Additional region of interest (ROI) analyses within the superior temporal plane were carried out in regions anatomically defined by the Jülich and Harvard-Oxford probabilistic atlases (distributed with FslView) and thresholded at 30%. These included primary auditory cortex (area Te1.0 in middle Heschl’s gyrus [HG]) and the non-primary auditory areas Te1.1 (posteromedial HG), Te1.2 (anterolateral HG), planum polare (PP) and planum temporale (PT). We also tested the posterior parietal region revealed in the whole-cortex SnPM analysis, to enable a comparison of effect size with the auditory cortical ROIs and to statistically test for between-region differences. To avoid statistical “double-dipping” (Kriegeskorte et al., 2009), we used a leave-one-subject-out procedure (Esterman et al., 2010) in which the whole-cortex second level t-test was repeatedly re-estimated, each time leaving out one participant, and using the resulting left parietal cluster as the ROI for the left out subject (cluster defining threshold p < .005 uncorrected). To obtain the homologous cluster in the right hemisphere, each left parietal cluster was left-right flipped using MarsBaR toolbox for SPM (http://marsbar.sourceforge.net). To reduce computation time, these leave-one-subject-out t-tests were conducted parametrically in SPM (i.e. without the SnPM toolbox). ROI effect sizes were computed by averaging the searchlight image over the spatial extent of each ROI. To facilitate interpretation (Allefeld and Haynes, 2014), ROI effect sizes are reported after transforming the standardized pattern distinctness back into the original estimate (by multiplying by a constant factor of √123 i.e. the typical number of voxels within each searchlight).

Classical multidimensional scaling (MDS) was performed on the average dissimilarity matrix in selected ROIs, formed by computing the pattern distinctness between all stimuli. Prior to MDS, each element in the dissimilarity matrices was subjected to a group-level one-sample t-test. Given that the goal of this analysis was to better visualize effects of interest already identified as significant (i.e. the independent and integrated contrasts in the whole-cortex and ROI analyses), we thresholded these dissimilarity matrices at p < .05 uncorrected.

### Spatial resolution of current fMRI data and relationship with previous mapping studies

Because we wished to measure whole-brain responses, including in regions outside classically defined auditory cortex, we measured BOLD responses with a resolution of 3 mm isotropic voxels (the data were additionally smoothed with a 6 mm kernel but only after the critical multivariate statistics were computed). While finer-resolution data are commonly obtained in studies investigating how frequency and other acoustic features are mapped to individual voxels (e.g. Formisano et al., 2003; Barton et al., 2012; Herdener et al., 2013; Leaver and Rauschecker, 2016), our concern here is how frequency and AM features are represented at a more abstract level in activity patterns over multiple voxels. Such representations may reflect both “distributed” and “sparse” coding schemes (Bizley and Cohen, 2013). It is well-established that multivoxel methods can sensitively measure changes in brain responses to acoustic features (even with standard-resolution data) by pooling weak but consistent signals over voxels and exploiting between-voxel correlations (e.g. Linke and Cusack, 2015).

Note that while significant independent coding of frequency and AM might be consistent with separate underlying neural populations responding to those features, this need not be the case. That is, the same neurons could simply be responding in a linear (additive) fashion to changes in frequency and AM rate. Thus, the extent of representational independence and integration in multivoxel patterns reveals more abstract computational properties (rather than the precise spatial configuration) of neural populations in a cortical region.

## Results

### Cortical distribution of independent and integrated codes

We used cross-validated MANOVA (Allefeld and Haynes, 2014) to determine the extent to which cortical activity patterns show evidence for 1) independent coding of frequency, in which the influence of frequency was invariant with respect to AM, 2) independent coding of AM, in which the influence of AM was invariant with frequency or 3) integrated coding, in which the influences of frequency and AM were interdependent. This was achieved by testing whether the pattern distinctness D over a searchlight sphere or ROI was significantly above zero for the independent and integrated effects of interest (see First-level statistics in the Methods section).

Using a whole-cortex searchlight analysis (Kriegeskorte et al., 2006), we detected large clusters in the superior temporal plane bilaterally (extending into the superior temporal gyrus) that showed significant independent coding of frequency and AM (Figure 2A and Table 1). Within these regions of auditory cortex, there was no evidence for integrated coding after correcting for multiple comparisons over the whole cortex. Instead, significant integrated coding was observed in a cluster outside of classically defined auditory cortex in the left posterior parietal lobe, extending over the inferior and superior portions of the parietal lobule and the intraparietal sulcus.

**Table 1.**
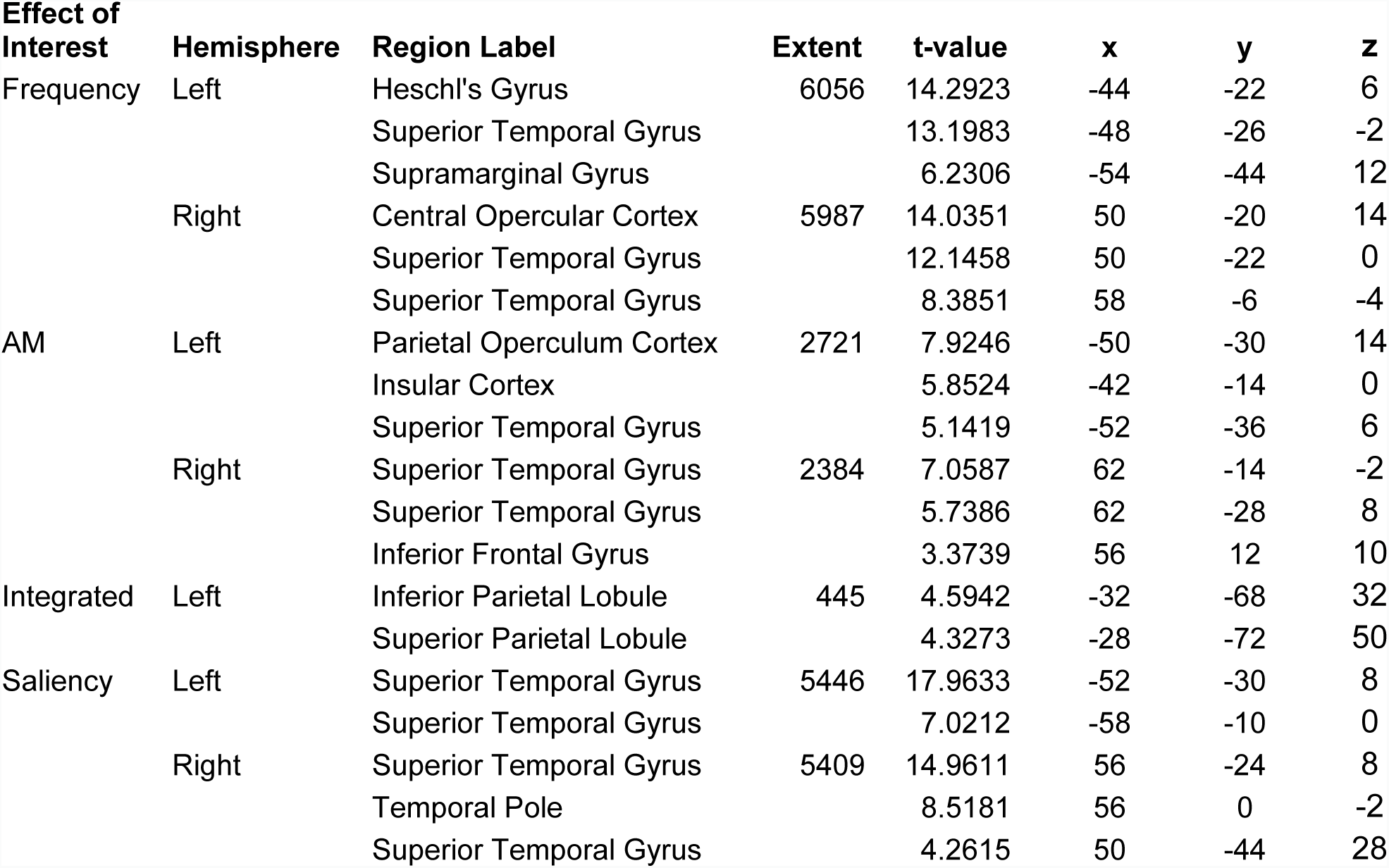
MNI coordinates and anatomical labels for significant multivariate searchlight effects

| Effect of Interest | Hemisphere | Region Label | Extent | t-value | x | y | z |
| --- | --- | --- | --- | --- | --- | --- | --- |
| Frequency | Left | Heschl's Gyrus | 6056 | 14.2923 | -44 | -22 | 6 |
|  |  | Superior Temporal Gyrus |  | 13.1983 | -48 | -26 | -2 |
|  |  | Supramarginal Gyrus |  | 6.2306 | -54 | -44 | 12 |
|  | Right | Central Opercular Cortex | 5987 | 14.0351 | 50 | -20 | 14 |
|  |  | Superior Temporal Gyrus |  | 12.1458 | 50 | -22 | 0 |
|  |  | Superior Temporal Gyrus |  | 8.3851 | 58 | -6 | -4 |
| AM | Left | Parietal Operculum Cortex | 2721 | 7.9246 | -50 | -30 | 14 |
|  |  | Insular Cortex |  | 5.8524 | -42 | -14 | 0 |
|  |  | Superior Temporal Gyrus |  | 5.1419 | -52 | -36 | 6 |
|  | Right | Superior Temporal Gyrus | 2384 | 7.0587 | 62 | -14 | -2 |
|  |  | Superior Temporal Gyrus |  | 5.7386 | 62 | -28 | 8 |
|  |  | Inferior Frontal Gyrus |  | 3.3739 | 56 | 12 | 10 |
| Integrated | Left | Inferior Parietal Lobule | 445 | 4.5942 | -32 | -68 | 32 |
|  |  | Superior Parietal Lobule |  | 4.3273 | -28 | -72 | 50 |
| Saliency | Left | Superior Temporal Gyrus | 5446 | 17.9633 | -52 | -30 | 8 |
|  |  | Superior Temporal Gyrus |  | 7.0212 | -58 | -10 | 0 |
|  | Right | Superior Temporal Gyrus | 5409 | 14.9611 | 56 | -24 | 8 |
|  |  | Temporal Pole |  | 8.5181 | 56 | 0 | -2 |
|  |  | Superior Temporal Gyrus |  | 4.2615 | 50 | -44 | 28 |

**Figure 2.**
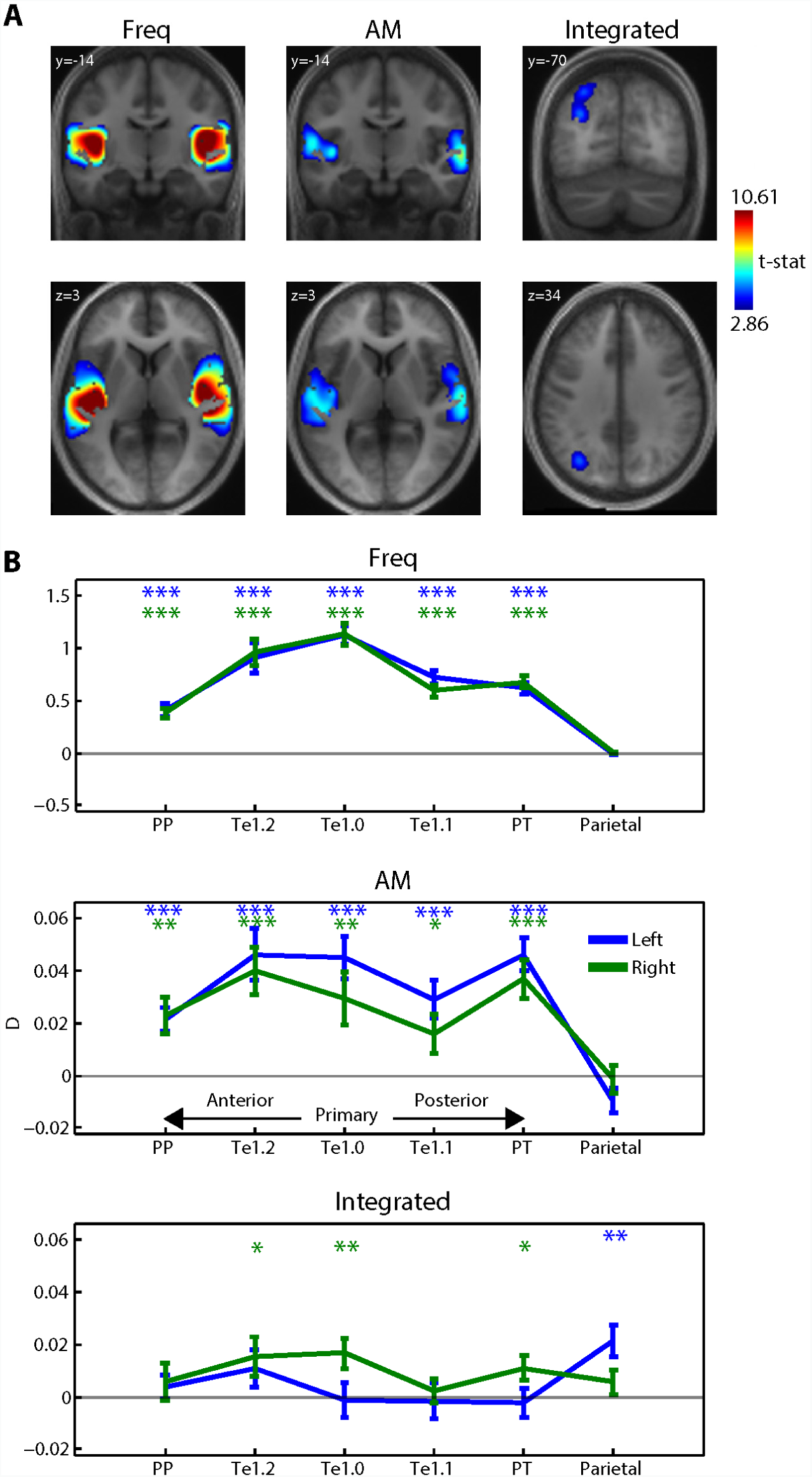
Whole-cortex multivariate searchlight analysis. A) Group-level statistical maps for each effect of interest, overlaid onto coronal and axial sections of the group-averaged structural (in MNI space) and thresholded voxelwise at p < .005 and clusterwise at p < .05 (FWE corrected for multiple comparisons). B) ROI analysis. Each data point shows the pattern distinctness D, averaged over the searchlight map within each ROI and over participants. Error bars represent the standard error of the mean. Asterisk symbols above each data point indicate significantly above-zero pattern distinctness, FDR corrected for multiple comparisons across contrasts, ROIs and hemispheres. *** p < .001, ** p < .01, * p < .05.

We next conducted an ROI analysis in which independent and integrated coding was tested in anatomically defined regions in the superior temporal plane, including primary auditory cortex in middle HG (area Te1.0) as well as regions more anterior (Te1.2, PP) and posterior (Te1.1 and PT). This allowed us to make between-region comparisons and examine how the strength of independent and integrated codes changes with increasing levels of the cortical hierarchy. In addition to the anatomically defined auditory ROIs, we included the posterior parietal region identified in the whole-cortex searchlight analysis. To avoid statistical “double-dipping” (Kriegeskorte et al., 2009), this parietal region was functionally defined using a leave-one-subject-out procedure (Esterman et al., 2010).

We first tested each ROI separately, using false discovery rate (FDR) correction for multiple comparisons across 6 ROIs x 2 hemispheres x 3 effects of interest (Genovese et al., 2002). As expected from the earlier whole-cortex analysis, significant independent coding of both frequency and AM was observed in all auditory ROIs but not in posterior parietal cortex (shown in Figure 2B). The effect size for independent coding of AM (mean D = 0.02-0.04 over auditory regions) was relatively small, amounting to no more than 8% of the frequency effect size (mean D = 0.5-1.0). Also expected was significant integrated coding in the left posterior parietal ROI. However, additional effects of integrated coding were observed in right primary auditory cortex (area Te1.0), right anterolateral auditory area Te1.2 and right PT. The effect size for integrated coding (mean D = 0.01-0.02 over right Te1.0, Te1.2, PT and left parietal) was considerably smaller than that for independent coding (50% of the AM effect size and no more than 4% of the frequency effect size). Thus, this ROI analysis suggests that in sub-fields of auditory cortex, cortical activation patterns show a mixture of components: a strong independent code and a weak integrated code. In contrast in parietal cortex, only an integrated code is present.

Pairwise comparisons between left and right hemispheres revealed only one significant effect: an increase in frequency coding in left versus right auditory area Te1.1 (two-tailed pairwise t(19) = 2.55, p < .025). However, this did not survive FDR correction for multiple comparisons across regions.

We next assessed how the magnitude of independent and integrated coding changed along successive stages of the cortical hierarchy. For independent coding of frequency, there was a significant decrease in pattern distinctness in non-primary versus primary auditory cortex (t(19) = −12.2, p < .001; region x hemisphere interaction: t(19) = .427, p = .674). This was also the case for parietal versus non-primary auditory cortex (t(19) = −11.8, p < .001; region x hemisphere interaction: t(19) = -.612, p = .548). The pattern was less clear-cut for independent coding of AM and integrated coding. Like the results for the frequency feature, there was a significant decrease in independent coding of AM in parietal versus non-primary auditory cortex (t(19) = −5.38, p < .001; region x hemisphere interaction: t(19) = −1.89, p = .075). However, the equivalent comparison for non-primary versus primary auditory cortex was not significant (t(19) = −1.21, p = .240; region x hemisphere interaction: t(19) = −1.04, p = .312). For integrated coding, there was a (left-lateralized) increase in parietal versus non-primary auditory cortex (left hemisphere: t(19) = 2.94, p < .01; right hemisphere: t(19) = −.539, p = .596; region x hemisphere interaction: t(19) = 2.72, p < .025). However, there was no significant difference between non-primary and primary auditory regions (t(19) = −0.797, p = .435; region x hemisphere interaction: t(19) = 1.67, p = .112). In summary, although there was a clear and fine-grained change across hierarchical levels in the strength of frequency coding (primary vs. non-primary auditory cortex, non-primary auditory vs. parietal cortex), such a change for AM and integrated coding was less fine-grained and only evident in the higher hierarchical levels (non-primary vs. parietal cortex).

Additional univariate analyses were conducted in which we assessed the strength of activation in each ROI using repeated measures ANOVA (with frequency and AM as factors). As shown in Figure 3, main effects of frequency and AM were present in auditory cortical regions but not in parietal cortex (FDR corrected as before, across 6 ROIs x 2 hemispheres x 3 effects of interest). No significant interaction between frequency and AM was observed in any of the regions tested (even with an uncorrected threshold). This suggests that the integrated coding effects revealed by cvMANOVA are inherently multivariate and arise from the pattern (and not strength) of multivoxel activity, a point to which we will return in the Discussion.

**Figure 3.**
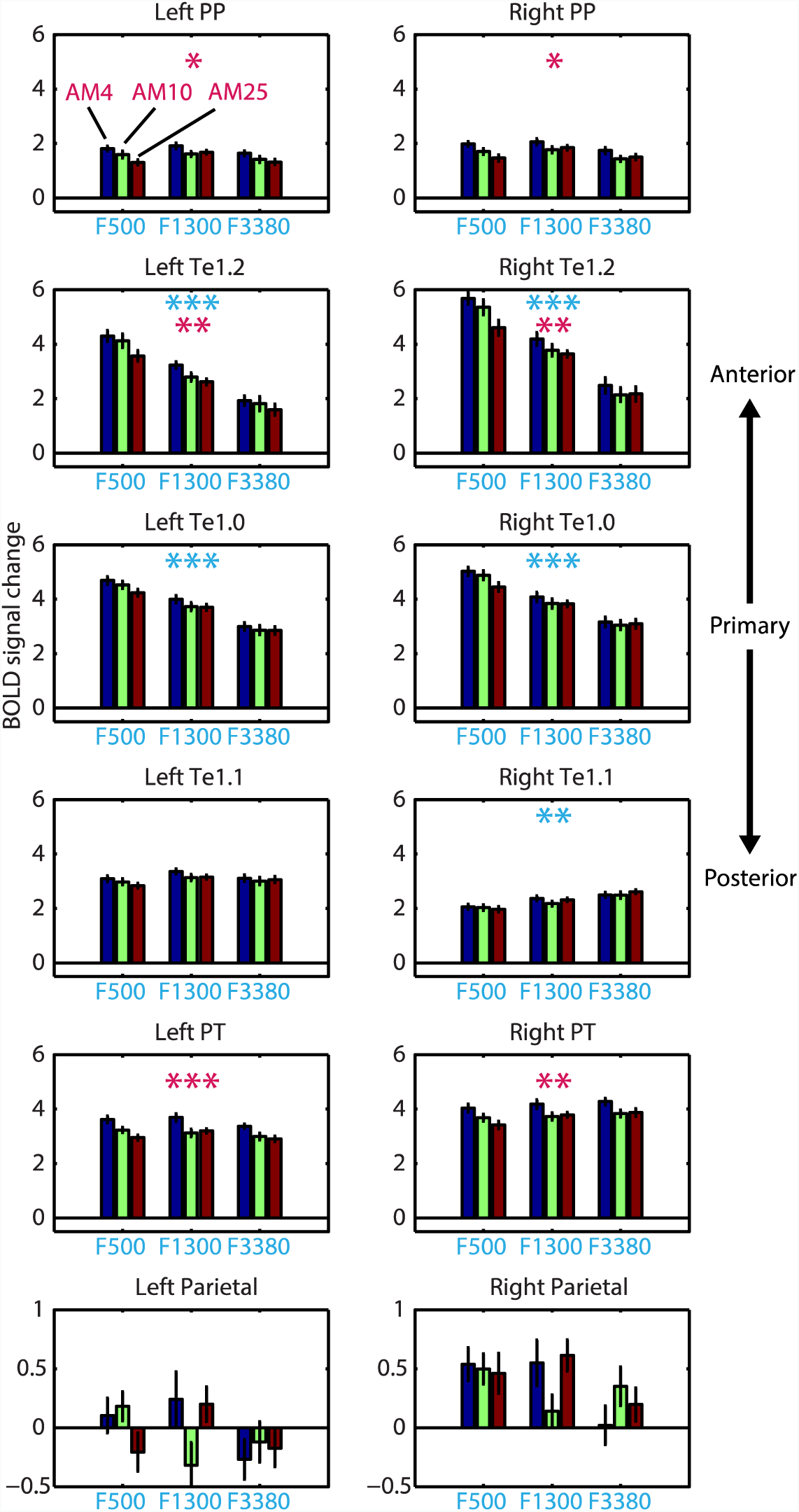
Univariate ROI analysis. Data represent the BOLD signal change averaged over the spatial extent of each ROI and across participants. Error bars represent the standard error of the mean. Asterisk symbols indicate a significant main effect of frequency (in cyan) or AM rate (in magenta), FDR corrected for multiple comparisons across contrast, ROI and hemisphere. *** p < .001, ** p < .01, * p < .05.

### Multidimensional scaling analysis

Having established the cortical distribution of independent and integrated codes, we next used classical MDS to further characterize those codes (Kriegeskorte and Kievit, 2013). In three selected ROIs (right Te1.0, right PT and left parietal), we computed the pattern distinctness for all pairs of stimuli and assembled the results into dissimilarity matrices. These ROIs were chosen as together they fully sample the transition from auditory core to non-core to parietal cortex and show a mixture of independent and integrated coding profiles. After averaging the matrices over participants and thresholding at p < .05 uncorrected (Figure 4A), MDS was performed to project the multivoxel dissimilarity structure onto a simple two-dimensional space (Figure 4B). In this visualization, stimuli that are close together are associated with similar multivoxel activation patterns while stimuli that are far from each other are associated with dissimilar patterns.

**Figure 4.**
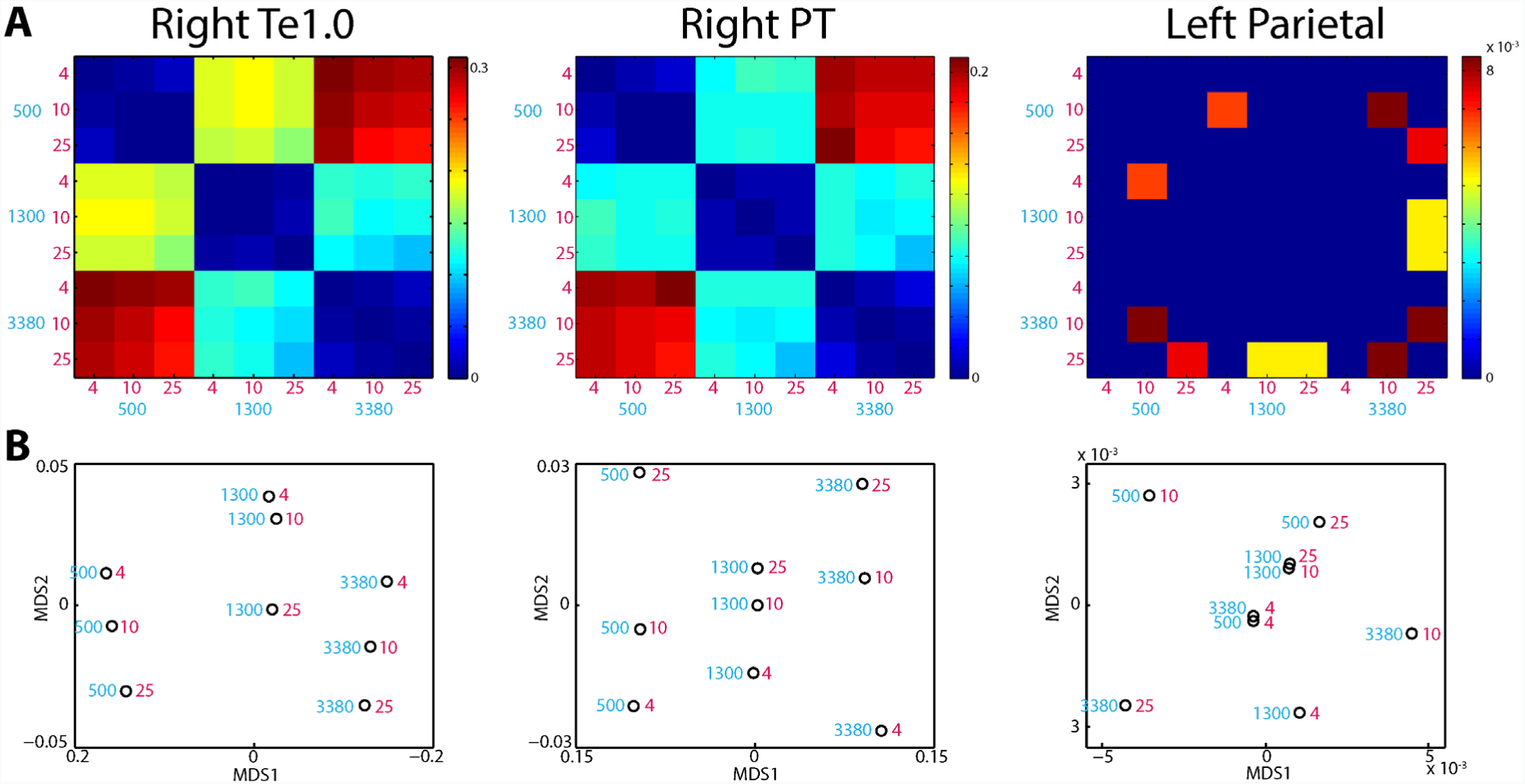
Visualizations of multivariate pattern distinctness A) Matrices expressing the multivoxel dissimilarity for all pairs of stimuli, averaged over the searchlight map within each ROI. Warm colors indicate multivoxel patterns that are highly dissimilar while cool colors indicate less dissimilarity. Dissimilarity matrices are shown thresholded at p < .05 (uncorrected). B) MDS solutions for the dissimilarity matrices shown in panel A (first two dimensions plotted only). The cyan number beside each data point indicates the carrier center frequency of the bandpass noise while the magenta number indicates the AM rate.

In right primary auditory cortex (area Te1.0) and right PT, frequency and AM features were automatically projected by the MDS solution onto separate dimensions, despite the method having no information as to the stimulus features. Frequency was carried by the first MDS dimension (shown as the x-axis in Figure 4B) while AM was carried by the second dimension (y-axis). This is consistent with our previous observation of these regions representing frequency and AM in a largely independent manner. We note further that the 4 and 10 Hz AM rates (in right Te1.0) and the 10 and 25 Hz AM rates (in right PT) were closer in MDS space for the middle carrier frequency (1300 Hz), which may account for the small degree of integrated coding observed in these regions. However, as establishing the group-level reliability of MDS solutions is difficult due to the arbitrary rotation induced by the method (Ejaz et al., 2015), we refrain from drawing strong conclusions about this latter observation.

In contrast to auditory cortex, MDS for the left parietal ROI did not clearly separate frequency and AM features. Instead, activation patterns in this region were modulated by particular conjunctions of carrier frequency and AM rate (e.g. F500AM10 and F3380AM25). This is again consistent with our previous observation that parietal cortex is characterized solely by an integrated code.

### Saliency analysis

In the visual domain, parietal cortex has repeatedly been implicated in the processing of bottom-up saliency (Arcizet et al., 2011; Bogler et al., 2011). We therefore asked to what extent the integrated coding effect observed in posterior parietal cortex could be explained by between-stimulus differences in perceived saliency. In a separate behavioral session, listeners listened to all pairwise combinations of the nine sounds and judged which sound in each pair was more salient. We then estimated the perceived saliency of each sound as the percentage of trials the sound was chosen as more salient (shown in Figure 5A as thick black line). Because saliency is related (although not identical) to loudness (Liao et al., 2015), we also show for comparison the loudness of the stimuli as predicted by the model of Moore et al. 2016 (shown in Figure 5A as thick blue line).

**Figure 5.**
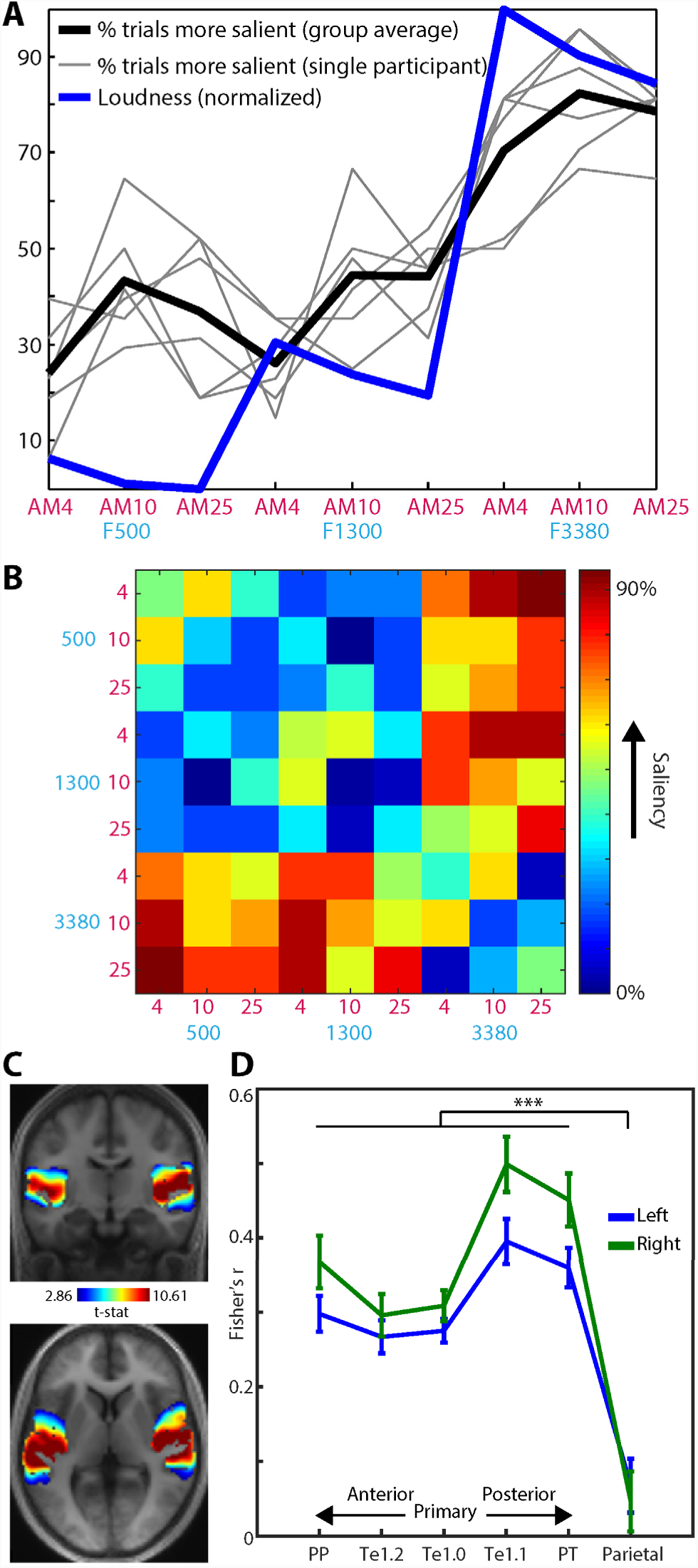
Saliency analysis. A) Subjective saliency of the stimuli. The thick black line indicates the group-averaged percentage of trials each stimulus was judged as more salient (than the other stimuli). Light gray lines indicate saliency judgements for individual participants. The thick blue line represents the predicted loudness of the stimuli according to the model of Moore et al. (2016) and normalized to have the same scale as the saliency data (for display purposes only). B) “Saliency distance” matrix expressing the absolute difference in the percentage of observations each sound in a pair was chosen as more salient. C) Whole-cortex multivariate searchlight analysis, showing where the Fisher transformed spearman correlation between the saliency distance matrix in panel B and the multivoxel dissimilarity structure in each searchlight was significantly above zero across participants (thresholded voxelwise at p < .005 and clusterwise at p < .05 FWE corrected for multiple comparisons). D) ROI analysis. Each data point shows the Fisher transformed Spearman correlation, averaged over the searchlight map within each ROI and over participants. Error bars represent the standard error of the mean. Brace and asterisk indicates significant p < .001 F-test comparing the strength of Spearman correlation between auditory and parietal regions.

Repeated measures ANOVA of the saliency judgments, with frequency and AM rate as factors, revealed a significant main effect of frequency (reflecting higher saliency for increasing frequency; F(2,10) = 31.5, p < .001) and a significant main effect of AM rate (reflecting higher saliency for the middle AM rate; F(2,10) = 6.34, p < .025). However, the interaction between frequency and AM rate was not significant (F(4,20) = .808, p = .512). To directly test whether there was positive evidence for the null effect of no interaction, we also conducted repeated measures ANOVA as a Bayesian analysis (Rouder et al., 2016, 2017; Marsman and Wagenmakers, 2017). We contrasted a model which contained both main effects of frequency and AM and their interaction, with a null model that had the same structure but lacked the interaction (both models were assigned a prior probability of 0.5). This analysis indicated that the null model was 5 times more likely than the alternative model (Bayes Factor = 5.31). As the integrated coding effect in parietal cortex is defined by the interaction between frequency and AM, the absence of an interaction in the saliency judgments is therefore inconsistent with a saliency-based account of the integrated coding effect in parietal cortex, or indeed, in any other of the regions in which integrated coding was observed.

As a further test of a saliency-based account, we used representational similarity analysis (RSA) to relate listeners’ saliency judgments to the observed multivoxel patterns (Kriegeskorte and Kievit, 2013). For each pair of sounds presented in the saliency judgment task, we pooled saliency judgments over trials and participants and computed the absolute difference in the percentage of observations each sound in the pair was chosen as more salient. From this we assembled a distance matrix quantifying the difference in saliency between the two sounds of all presented pairs (Figure 5B). This “saliency distance” matrix provides a more detailed characterization of between-stimulus differences in saliency than the summary measure presented in Figure 5A, which we could then correlate with the multivoxel dissimilarity matrix observed in each searchlight across the cortex of individual participants. As shown in Figure 5C, the (Fisher-transformed) Spearman correlation between the saliency and multivoxel dissimilarity structure was significantly above zero in the superior temporal plane bilaterally but not in parietal cortex (for MNI coordinates, see Table 1). This pattern was further supported by an ROI analysis (Figure 5D) in which the Spearman correlation significantly decreased from superior temporal to parietal cortex (F(1,19) = 57.8, p < .001; effects involving hemisphere were not significant). We further note with interest how this saliency-to-multivoxel correlation peaked in posteromedial auditory area Te1.1, which clearly differs to how the independent and integrated coding effects were expressed over cortical regions (compare Figure 5D with Figure 2B). Nearly identical results were obtained when using loudness in this ROI analysis (here a loudness distance matrix was formed by computing the absolute differences in loudness between the stimuli). This suggests that saliency/loudness can be reliably dissociated from the independent and integrated coding effects of the earlier analyses. In summary then, this RSA analysis together with the absence of interactive influences of frequency and AM on behavioral saliency judgments suggests that the integrated coding effect we observe cannot be attributed to saliency/loudness. We will return to this point in the Discussion.

## Discussion

In the current study, we manipulated two important acoustic features in parallel, frequency and AM rate, and determined the extent to which they are represented by independent versus integrated codes in fMRI multivoxel patterns. We demonstrate that these spectral and temporal dimensions are represented largely independently in the superior temporal plane, with only a weakly integrated component present in right Te1.0, Te1.2 and PT (amounting to no more than 4% of the frequency effect size and 50% of the AM rate effect size). In contrast, in a posterior parietal region not classically considered part of auditory cortex, neural representation is exclusively integrated albeit weakly.

### Independent representations in the superior temporal plane

Our demonstration of largely independent representations of frequency and AM rate in the superior temporal plane contrasts with evidence from animal physiology that suggest highly non-linear representations already at the level of primary auditory cortex (e.g. deCharms et al., 1998; Nelken et al., 2003; Wang et al., 2005). While there are many differences between the current study and this previous work (most obviously, species and recording technique), our findings may also reflect the specific features that were manipulated. Specifically, it has been suggested that frequency and AM rate are fundamental dimensions of sound analysis (Dau et al., 1997; Chi et al., 2005) and in the auditory cortex are represented as orthogonally-organized topographic maps (“tonotopy” and “periodotopy”; e.g. Baumann et al., 2015). Our findings in the superior temporal plane are thus consistent with the notion of orthogonal maps for frequency and AM features. While previous electrophysiological (Langner et al., 2009) and fMRI (Baumann et al., 2015) findings from animals also support this proposal, in humans the evidence is mixed with some studies showing clear topographic organization (Langner et al., 1997; Barton et al., 2012; Herdener et al., 2013) but others not (Giraud et al., 2000; Schönwiesner and Zatorre, 2009; Overath et al., 2012; Leaver and Rauschecker, 2016). These conflicting findings may be attributed to the small size of auditory cortex and high inter-subject variability in anatomy. In the current study we overcame these challenges by using a multivariate analysis method that abstracts away from the precise configuration of voxels. Importantly, this approach allowed us to directly test and quantify the degree of representational independence, an approach distinct to the more qualitative inferences of previous mapping studies.

Orthogonal representation of frequency and AM features is also suggested by component analysis of human fMRI responses to natural sounds (Norman-Haignere et al., 2015). This work suggests that frequency and AM features are represented as independent components in partly overlapping regions of the superior temporal plane. However, this study did not test for feature interactions between those features, leaving unclear the relative contributions of independent and integrated representations to neural responses.

Thus, our study provides new evidence that frequency and AM are orthogonal dimensions of sound analysis. Such independent representation may support listeners’ ability to selectively process information in frequency versus time. In addition, as noted by Schnupp (2001), an independent coding scheme will tend to convey more information than a highly-selective integrated code. This property would be desirable if the role of primary auditory cortex was to relay information to more specialized feature conjunction detectors in higher-level regions.

### Integrated representation in posterior parietal cortex

Our imaging of the entire cortex allowed us to probe beyond classically defined auditory cortex. In this respect, a striking demonstration here is of an exclusively integrated representation of frequency and AM rate in a left parietal region, just posterior to the intraparietal sulcus (IPS). This finding is notable for two reasons. First, it parallels findings from the visual domain in which parietal cortex (in particular the IPS) shows increased fMRI responses in feature conjunction versus single feature tasks (Donner et al., 2002; Shafritz et al., 2002; see also Baumgartner et al., 2013 for a similar finding using multivariate methods), with damage to this region leading to feature binding deficits (Humphreys et al., 2000). Second, BOLD activation in the IPS has been shown to systematically vary in auditory bi-stability (Cusack, 2005) and figure-ground paradigms (Teki et al., 2011, 2016). Indeed, the peak locations of the posterior parietal effects reported by these latter studies fall inside the cluster reported here. In both these auditory paradigms, perceptual outcomes are critically dependent on the way in which information across multiple features (frequency and time) is combined. Thus, the integrated representation for frequency and AM we observe here in posterior parietal cortex is consistent with previous work suggesting a role for the IPS in feature integration and the structuring of acoustic input, possibly alongside other parietal regions specialized for visual information (for further discussion, see Cusack, 2005). However, our study goes beyond previous work that measured overall regional differences in fMRI or MEG signal amplitude by more directly probing representational content in multivoxel patterns, an approach which is less susceptible to confounding factors such as task difficulty (Baumgartner et al., 2013).

Because of previous findings from the visual domain implicating parietal cortex in bottom-up saliency (Arcizet et al., 2011; Bogler et al., 2011), we also asked a separate group of listeners to rate the subjective saliency of the stimuli. While the sounds clearly differed in their subjective saliency, we found that influences of frequency and AM on the saliency ratings combined independently without evidence for an interaction, an observation inconsistent with a saliency based account. Moreover, when using RSA to relate saliency judgments to the dissimilarity structure of the multivoxel patterns, we found that saliency did not correlate with multivoxel patterns in parietal cortex. Rather, the effect of saliency was confined to superior temporal plane regions with a peak in posteromedial auditory area Te1.1, which is reminiscent of findings by Behler and Uppenkamp (2016) who reported correlates of loudness in this region (see Liao et al., 2015 for the close relationship between loudness and saliency). Thus, the results from this saliency analysis suggest that the integrated coding effect we observe cannot be attributed to bottom-up saliency.

Related to the issue of saliency, we also consider the possibility that the integrated coding profile we observe in parietal cortex was in part a consequence of listeners’ task. In our study, listeners performed an attentionally undemanding task that did not require explicit integration of frequency and AM features: detecting the target white-noise interruptions could in principle be based on changes in either the amplitude or spectral profiles alone. Despite this, one might argue that participants nevertheless detected the noise interruptions by attending to changes in both temporal and spectral content, in turn contributing to the integrated coding effect we observe. Indeed, as discussed below, attention has long been proposed to mediate feature integration (Treisman and Gelade, 1980). However, we think that this is unlikely as an explanation for the current findings. The interaction between frequency and AM rate in parietal cortex resulted from differences in the multivoxel patterns evoked by our stimuli (while the task was fixed throughout). Thus, even if listeners monitored both spectral and temporal content to detect the target interruptions, it is unclear how this would have preferentially biased listeners’ attention towards certain feature conjunctions. This is because the targets were temporally unmodulated and spectrally wide-band and therefore “neutral” with respect to the nine feature conjunctions of the stimuli.

A key assumption in our approach to distinguishing independent and integrated representations is a linear relationship between underlying neural activity and the measured fMRI signal (Kornysheva and Diedrichsen, 2014; Erez et al., 2015). Our univariate analysis shows that the mean signal amplitude in the posterior parietal region did not differ between stimuli, neither in terms of mains effects nor in the interaction between frequency and AM rate. This suggests that our experimental manipulations in this region did not evoke sufficiently large changes in mean signal to saturate the fMRI response and produce non-linear signal changes that could be misinterpreted as an integrated representation.

The integration of multiple feature representations is critical for building a cohesive perception of the auditory scene. However, even in parietal cortex, the effect size for integrated coding was small in comparison with that observed for independent coding in the superior temporal plane. Why then do we observe only weak integration of frequency and AM rate? As discussed above, frequency and AM may be privileged dimensions of sound analysis that are separable in a way that other dimensions are not. Our results may also be attributed to listeners performing an attentionally undemanding task that did not require explicit integration of frequency and AM features. It has been suggested that while individual features are detected automatically, feature integration is a computationally demanding process requiring focused attention (Treisman and Gelade, 1980; Shamma et al., 2011). Thus, the absence of focused attention to feature conjunctions could explain the weak integration we observe. Future work, using manipulations of attention, will be required to test this proposal.

## Acknowledgements

We are grateful to Carsten Allefeld for advice with crossvalidated MANOVA.

